# Chemodiverse cell systems responses to UV in an algal sister of land plants

**DOI:** 10.64898/2026.02.10.704744

**Authors:** Cäcilia F. Kunz, Ilka N Abreu, Tatyana Darienko, Janine Fürst-Jansen, Kirstin Feussner, Ivo Feussner, Maike Lorenz, Jan de Vries

## Abstract

Plant terrestrialization necessitated that a barrage of stressors had to be overcome^1^. Land plants use an integrated response network in adjusting their molecular physiology to terrestrial stressors^2^—one of the foremost being UV irradiance. The zygnematophytes are the closest streptophyte algal relatives of land plants^3-5^, are renowned for their resilience to UV stress^6-8^, and thus allow to glean key information for inferring the UV response toolkit of the earliest land plants^9,10^. Throughout streptophyte evolution, specialised metabolism radiated into creating diverse compounds used for responses to environmental challenges, such as sun-shielding compounds and antioxidants^11-14^. This includes UV-shielding compounds like flavonoids and coumarins but also the land plant specific polymer lignin, giving structural support in vascular plants^15^; homologs of the underpinning core pathway occur in streptophyte algae^16^. Here, we exposed the zygnematophyte *Mesotaenium* to UV-B irradiation and profiled its physiology, morphology, transcriptomics as well as metabolomic features. After UV-B exposure, cells showed rapid photophysiological responses and progressively growing terminal vacuoles. Our transcriptome data capture dynamic changes in gene expression of (i) core downstream responses such as genes homologous to phenol metabolic enzymes, photophysiological homeostats, and DNA repair factors; but also (ii) upstream components featuring key homologs of kinase-mediated signalling cascades, as well as light quality and abscisic acid-mediated signalling components. To scrutinize the acclimatory chassis, we created a metabolite feature database specifically for the *Mesotaenium* metabolome. Upon UV-B exposure, the metabolome displayed pronounced temporal shifts, with several phenolic features that accumulate along the stress–acclimation kinetics. Overall, we capture a chemodiverse response including various phenolics such as purpurogallin-like, methoxypsoralen-like derivatives and coumarins. Our data establish an integrated model for UV responses in the closest algal relatives of land plants, shedding light on the toolkit that allowed the progenitors of land plants to move out of a protective water column.

## Results and Discussion

### Dynamic physiological changes induced by UV exposure in an algal sister to land plants

Embryophytes derive from a single ancestor that had conquered land. For terrestrialization to succeed, many abiotic and biotic challenges must have been overcome, requiring that the earliest land plants dynamically adjusted to rapidly changing environmental conditions^1^. We worked with the unicellular genome-sequenced^17,18^ zygnematophyte *Mesotaenium endlicherianum* SAG 12.97, one of the closest algal relatives of land plants (Figure 1A). The algae were grown at 80 - 90 µmol photons m^-2^ s^-1^ for 7 days. 24 h before experimental start they were transferred to the experimental set-up. They were subjected to either the 50±10 µmol photons m^-2^ s^-1^ (Figure 1B). Upon experimental start, UV specimens were exposed to 3 hours of 1.7 - 2.9 W m^−2^ of ultraviolet (UV) radiation with a peak in the UV-B spectrum (at 311 nm, Figure S1A). To assess the gross physiological effects of UV-B on the algae, we tracked the photosystem II maximum efficiency (Fv’/Fm’) during the experiment (Figure 1C). Within the first hour, the average Fv’/Fm’ dropped from 0.525±0.027 to 0.289±0.047 (Wilcoxon test, Benjamini-Hochberg adjusted *P* = 0.022). This signifies that UV-B had a fast impact on the physiology. During recovery, the Fv’/Fm’ remained at lower levels of 0.328±0.0286 for UV versus 0.482±0.0209 for control (*P* = 6.41e-05), likely reflecting both the rapid response to UV and the slow and progressive acclimatory process. During the recovery phase both the acclimatory process but also perturbations that occurred during treatment and continue to effect the cells bears out; this aligns both with the accumulation of flavonoids—to an even higher degree than during the treatment itself—during the recovery of UV-treated *Marchantia*^19^ as well as lower Fv’/Fm’ in recovering *Zygnema* exposed to desiccation stress^20^.

**Figure 1.**
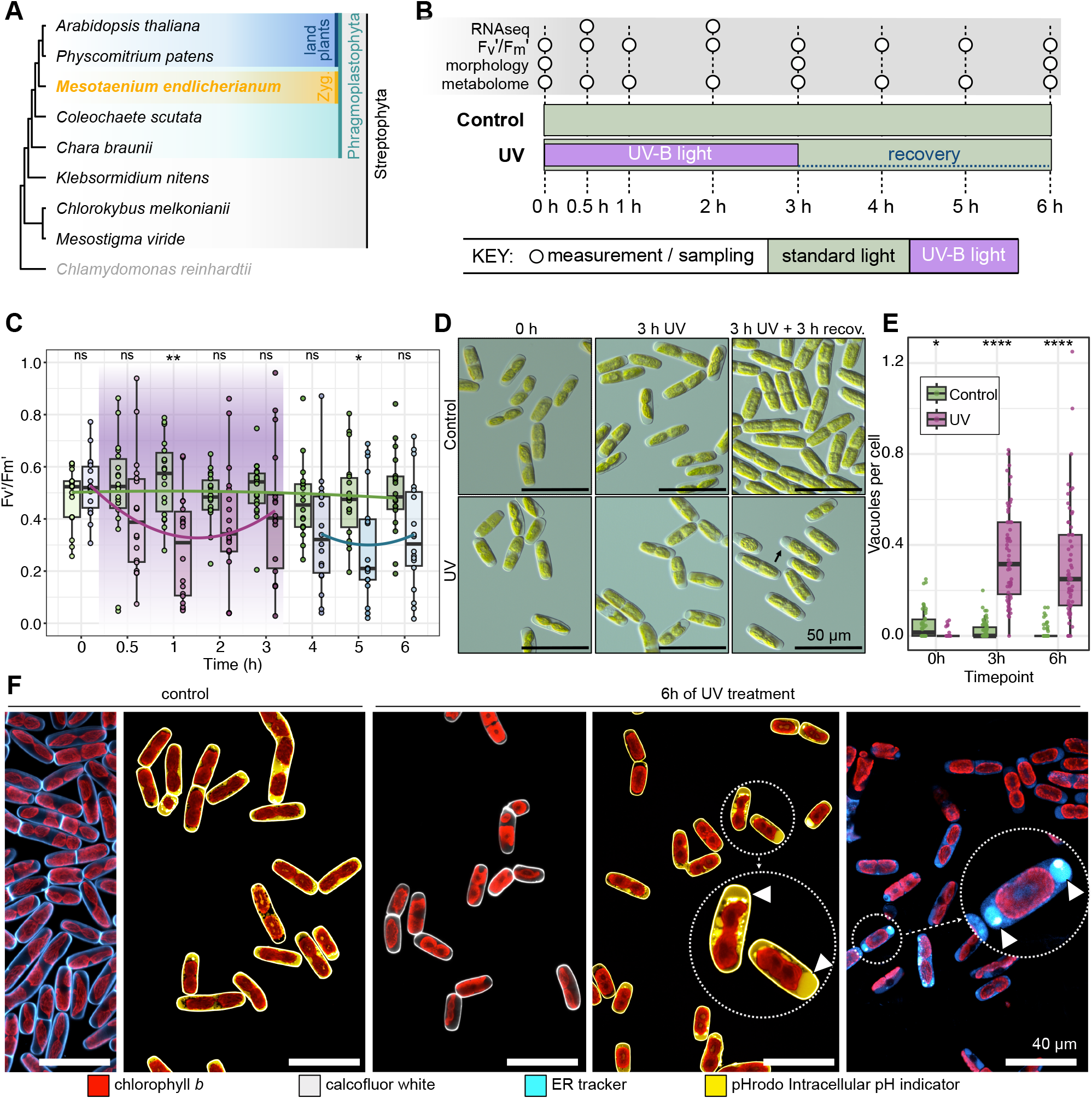
Rapid and quantifiable impact of UV on zygnematophyte cell physiology. (A) Cladogram of Streptophyta (grey) and a chlorophyte, indicating phragmoplastophyta (green), land plants (blue) and the here studied zygnematophyte (yellow). (B) Timeline of UV-B Experiment UV-B irradiation (purple) of 1 - 2.3 W m^−2^ was applied for 3 h in addition to photosynthetically active radiation (PAR; green) facilitated by grow lights with an intensity of 40 - 60 μmol m^−2^ s^−1^ followed by 3 h without UV-B irradiation. The sampling and measurement time points are indicated by circles (top) with a timeline (bottom), RNAseq: transcriptomics samples; Fv’/Fm’: measurements of PSII max. efficiency; morphology: microscopy images; metabolome: samples analyzed by untargeted UHPLC QTOF MS. (C) Photosystem II (PSII) maximum efficiency (Fv’/Fm’) at each timepoint. Control measurements of Fv’/Fm’ indicated in green, measurements of UV exposed plates indicated in purple (after UV-B exposure) to blue (after UV-B exposure ceased). Purple background indicates the Fv’/Fm’ measurements that were taken when UV-B irradiation was turned on. Significance levels depicted on top, pairwise comparison of each UV-B measurement with its corresponding control. Quadratic polynomial trend lines were fitted to each treatment phase (control, UV stress, UV recovery) using least-squares regression. Measurements taken with WALZ Mini PAM. (D) Control in top row, UV-B exposed samples in bottom row. First column corresponds to 0 h, second column to 3 h, and third column to t_6h_ (3 h UV treatment followed by 3 h of recovery). Scale indicated in bottom left corner of each image. UV 6 h contains an enlarged image of a cell, with an arrow pointing to a vacuole filled with quickly moving particles. (E) Boxplot that depicts the quantification of vacuole abundance (F) Confocal microscope panel of control and 6 h continuous UV-B exposure. Chlorophyll autofluorescence (red) was used as well as stains for cell wall (calcofluor white, grey), ER tracker (blue) and pHrodo intracellular pH indicator (yellow). Scale indicated in bottom right corner of each image. The two rightmost images contain enlarged images with arrows pointing at vacuoles.

*Mesotaenium endlicherianum* belongs to the order Serritaeniales^5^ within the Zygnematophyceae. Species of the genus *Serritaenia* form extracellular mucilage with UV protection based on compounds of unknown identity^21^. For microalgae, the intracellular accumulation of sunscreens is more common^21^. In *Mesotaenium endlicherianum*, the formation of extracellular mucilage as a response to environmental stress has been reported^22^, yet without accumulation of UV protective function. We therefore monitored the intracellular and extracellular responses to UV (Figure 1D). We noticed that the number of terminal vacuoles increased significantly by 15.4-fold at 3 h UV treatment and by 20.0-fold after subsequent 3 h of recovery as compared to control (P_adj_ = 1.35e-23 and 4.95e-18, respectively; Figure 1D,E). To further scrutinize the nature of these vacuoles, we used fluorescent stains for labelling intracellular pH changes (pHrodo) and an endoplasmic reticulum (ER) stain (ERtracker) to specifically observe membrane-bound acidic subcellular compartments via confocal laser scanning microscopy (cLSM; Figure 1F). After 3 h and 6 h (confocal images only) of UV treatment, defined terminal vacuoles become visible, discernable based on the ER tracker and the pH indicator (Figure 1F). Additionally, cell and chloroplast shapes appeared huddled (Figure 1F). This might be a mechanism of chloroplast movement to avoid excess light absorption^23^. The observed retraction of chloroplasts is in concordance with similar observations on the filamentous zygnematophytes *Spirogyra* and *Mougeotia* under heat stress^24^ and on diatoms, where UV-B exposure triggers decreases in chloroplast volume^25^.

Overall, there are defined and rapid cell physiological responses in our model system for the closest algal relatives of land plants. We next turned to understanding how this is reflected on the molecular level.

### The molecular response to UV in *Mesotaenium* builds on a conserved stress chassis

Zygnematophytes share key components of the molecular chassis for stress response with land plants, including high light, osmotic, and temperature stress^18,20,26-29^. To understand the impact of UV-B stress on the molecular physiology of the zygnematophyte *Mesotaenium*, we performed time course RNAseq analyses. We sequenced biological triplicates of two stressed and two control conditions (0.5 h and 2 h timepoints) on the illumina NextSeq platform. We obtained a total of 962,260,484 paired-end reads, cumulating to 144.4 Gbp of data. Using kallisto^30^, we pseudoaligned the reads onto the V2 gene models^18^ of the *Mesotaenium* genome^17^ at a rate of 85.89±2.24%. The gross transcriptomic profiles separated already after 0.5 h into treated and non-treated samples, a trend that increased with progression of treatment (Figure 2A, Figure S2A); the biological triplicates for each time and condition clustered together. We traced the progressive effect of UV treatment on the expression patterns of individual genes (Figure 2B). 241 genes were differentially regulated after 0.5 h (log_2_FC ≥ 1) and showed a ∼2.4-fold differential increase in transcript levels at 2 h (Figure S2B).

**Figure 2.**
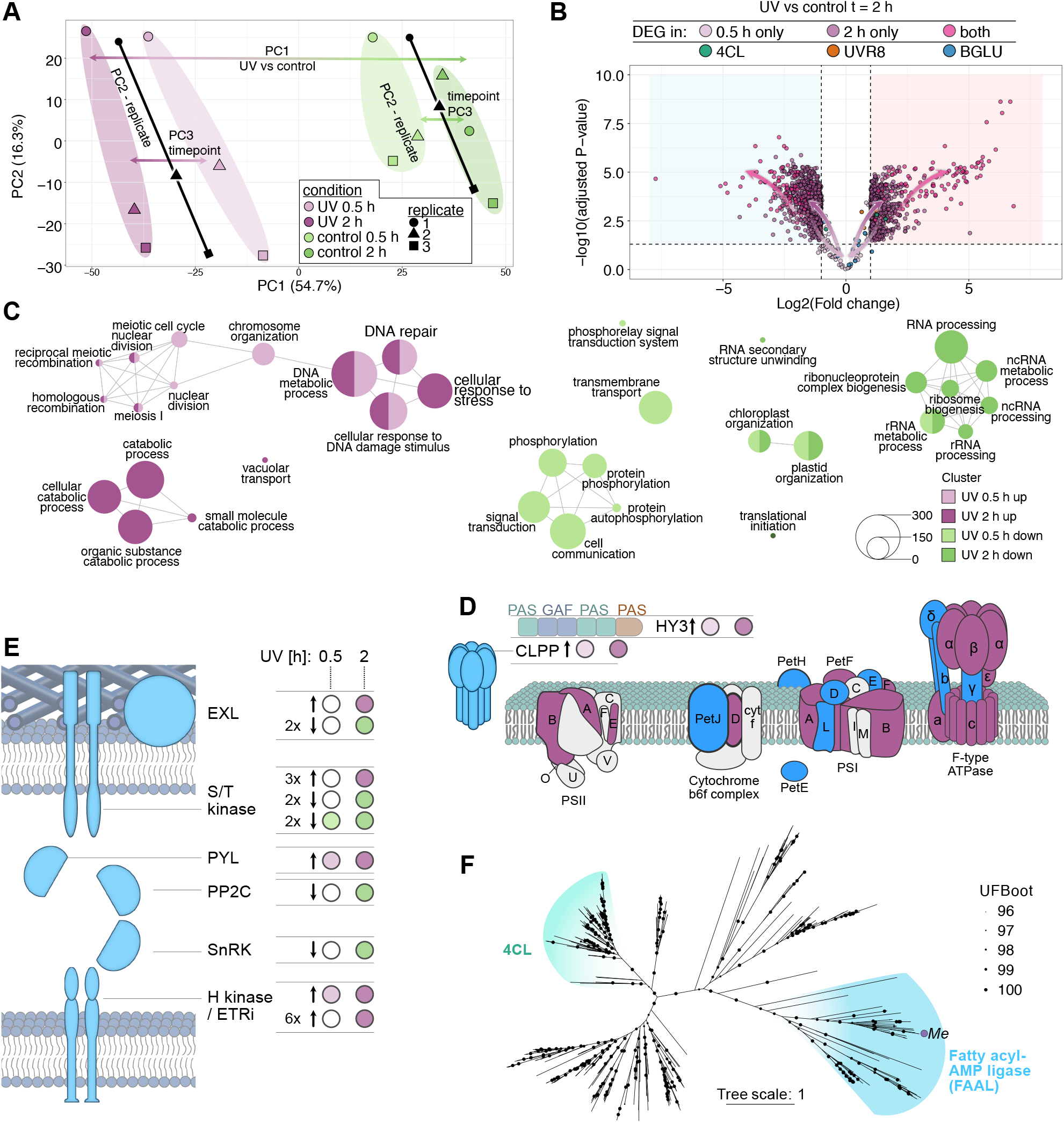
UV induces global transcriptional changes in genes for upstream and downstream acclimation. (A) Principal Component Analysis (PCA) of the filtered and normalised transcriptomic timepoints, indicating UV samples (pink) and control samples (green). Each timepoint has a specific shade of colour indicated on the right. The samples organise themselves according to condition and their timepoint from the middle to the outside. PC2 describes the variance of the samples which relates to the biological replicate. (B) Volcano plot of UV-B versus control samples at 2 h. x-axis: log2(FC) where Fold Change (FC) = |1.0|; y-axis: -log10(p) where the t-test p-value threshold is 0.05. Light purple indicates transcripts only differentially regulated at 0.5 h, pink in both 0.5 h and 2 h and dark purple only in 2 h. Arrows describe the change of transcripts through time. Green indicates homologs of 4CL, orange of UVR8, blue of BGLUs. Blue shading indicates downregulated, pink shading upregulated genes. (C) GO terms upregulated at 0.5 h (light purple) and 2 h (dark purple) and downregulated at 0.5 h (light green) and 2 h (dark green). (D) Protein domain structure (Per/Arnt/Sim, PAS; cGMP-specific phosphodiesterases, adenylyl cyclases and FhlA, GAF) of HY3 homolog and its upregulation after both 0.5 h and 2 h. Schematic of photosynthesis proteins, homologous transcripts for which were upregulated (purple) and present (blue). (E) Potential molecular circuitry of EXORDIUM-like proteins (EXL), Ser/Thr kinases, PYR1-like (PYL), PROTEIN PHOSPHATASE TYPE 2C (PP2C), SNF1-RELATED PROTEIN KINASE (SnRK), HISTIDINE KINASE (HK) / ETHYLENE RECEPTOR-like (ETRi) and number of homologs and their differential regulation after 0.5 h (light purple/green) and 2 h (dark purple/green). (F) Maximum likelihood phylogeny of 4CL homologs; the homolog of *Mesotaenium endlicherianum* (Me) that is most upregulated is indicated, clustering within a clade for fatty acyl-AMP ligases (FAAL) and not the 4CL group. The tree is drawn to scale (expected substitutions per site); UFBOOT values are indicated as scaled dots.

On balance, the data captured large parts of the molecular chassis of what has emerged as the stress response program that zygnematophytes share with land plants^18,26,27,31^; this chassis describes the logic of the network that underpins molecular physiological adaptation—from environmental cue input (here UV stress) to downstream output. In the following, we will move along that logic.

The key UV response indicator, the conserved transcription factor *UV-B RESISTANCE 8 (UVR8)*^32^ that acts as UV response hub^33,34^, was upregulated by 2.4-fold at 2 h of UV treatment (Figure 2B). In addition to UV specific photoreceptors, cryptochromes as well as phytochromes were upregulated. We found a homolog of the phytochrome-coding gene *LONG HYPOCOTYL 3* (*HY3*) to be upregulated 3.1-fold and 7.8-fold at 0.5 and 2 h after onset of UV exposure, respectively (Figure 2D). This signifies a rapid perception of the change in light quality and aligns with previous findings on another Serritaeniales, *Serritaenia testaceovaginata* ^35^. Phytochromes are among the key players for light quality signaling^36,37^, linking intracellular signalling and plastidial biology.

The chloroplast is recognized as an intracellular sensory compartment and the first frontier in stress response^38,39^. Indeed, among the pronouncedly induced genes—next to those coding for very short proteins—were those coding for components of photosystem I PsaA and PsaB and photosystem II PsbB PsbD (Fig. 2D). Further homologs of the important chloroplast protein homeostasis regulators Caseinolytic Protease Proteolytic subunit (CLPP)^40-42^ were upregulated 9.4- and 22.8-fold after 0.5 and 2 h of UV, respectively; diverse CLPP homologs also appeared as a stress hub in *Mesotaenium* co-expression modules correlated with high light^18^ (Fig. 2D). Apart from these upregulated homologous genes involved in chloroplast biology, GO term enrichment showed chloroplast organization to be broadly downregulated. It also showed DNA damage to be upregulated, among which homologs to genes involved in UV specific repair mechanisms like nucleotide and base excision repair such as TFIIH3, XPG, NTH, TDP1 and PCNA were involved (Figure 2C).

Zygnematophytes stand out from all other algae by having a full genetic chassis homologous to the canonical abscisic acid (ABA) signaling cascade^17,43^. While this deeply conserved cascade is likely acting ABA independently^44,45^, its components co-express since the last common ancestor (LCA) of zygnematophytes and land plants^31^. We find PYRABACTIN RESISTANCE-LIKE (PYL) upregulated by 5.7-fold while all ABA-relevant TYPE 2C PROTEIN PHOSPHATASES (PP2C) and SNF1-RELATED PROTEIN KINASE (SnRK) were downregulated (Fig. 2E). This could suggest a combination of a feedback loop and a phosphorelay shift to His and Ser/Thr kinases.

Recently, a track of kinase-coding genes was described to form the backbone of hubs in a predicted gene regulatory network shared by zygnematophytes (including diverse data derived from *Mesotaenium*) and land plants^27^. Here, under UV stress, several AHKs (Arabidopsis HISTIDINE KINASE) that were previously highlighted^27^ were significantly upregulated after 2 h and one even from 0.5 h onwards (Figure 2E). AHKs such as the ethylene-receptor-related histidine kinases (ETR-HKs) are not only sensing ethylene signalling, but a group of ETR-HKs was additionally found to be essential for ABA and osmostress signalling in *Physcomitrium patens*^46^. Disruption of ETR-HKs prevents the ABA-dependent autophosphorylation of RAF-like kinase. Therefore, the interaction of HK with RAF-like kinase can serve as an integration unit linking ethylene signalling and ABA/osmostress signalling pathways^46^. The Ser/Thr kinases, to which the RAF-like kinase belongs to, have been identified as the single most connected receiver in a predicted deep evolutionary gene regulatory network, therefore serving as a crucial hub of stress response^27^. Connected to other conserved hubs, the downstream responses to environmental inputs can be mediated, changing growth and acclimation. Thus, genes coding for parts of the kinome that represent major conserved proteins at the cell surface (in multicellular systems acting also in cell–cell communication) as well as intracellular signalling were pinpointed by our data; these likely formed nexuses in the UV response toolkit shared by land plants and their closest algal relatives.

We next turned to understanding how this translates into downstream molecular physiological acclimation. Upon UV, this prominently features downstream output on growth and physiology such as *EXORDIUM LIKE* (*EXL*) homologs, which in *Arabidopsis* regulate cell wall expansion during growth and stress^47,48^ and featured prominently in both transcriptomic^18,27^ and proteomic^49^ data of zygnematophytes exposed to diverse stressors.

Phenolics are key in UV response^50,51^. Our transcriptome data capture the expressional changes in diverse enzymes salient to specialized metabolism, including the hyperdiverse CYP450 which contributed greatly to the radiated chemodiversity of embryophytes^52-55^. Among the early and time-progressing responses were homologs of 4CL, and multiple β-glucosidases (BGLUs) (Figure 2B). BGLUs are stress induced and some can lead to the activation of coumarins via hydrolysis from their previous inactive storage form as glycosides^56,57^. In previous analyses the 4CL homologs of algae clustered co-orthologous to the *bona fide* 4CLs and other enzymes of land plants^58^. We analyzed the phylogeny of the 4CL homolog most up-regulated in Mesotaenium and found that it placed within a clade of fatty acyl-AMP ligase (FAAL) (Figure 2F). It belongs to the class I adenylate forming enzyme that have been suggested to act in polyketide biosynthesis^59,60^. We thus next honed in on dissecting the specialized metabolic profile of *Mesotaenium*.

### *Mesotaenium* exhibits a chemodiverse response to UV

It is a standing question in the field to what degree streptophyte algae utilize the phenylpropanoid pathway^58,61-64^. At the same time, it is known that different classes of streptophyte algae produce lineage-specific phenolics that act under stress^6,7,21,65^. We therefore used a time-course untargeted metabolomics approach to understand the chemodiversity with a particular focus on the repertoire of phenolics produced under UV-B exposure in *Mesotaenium*. For a time-course of t = 0 h, 0.5 h, 1 h, 2 h, 3 h, 4 h (3 h UV + 1 h recovery), 5 h (3 h UV + 2 h recovery), and 6 h (3 h UV + 3 h recovery), sampled in biological and technical triplicates, we obtained in total 1478 features based on m/z signals. These are grouped by time and treatment (Figure 3A).

**Figure 3.**
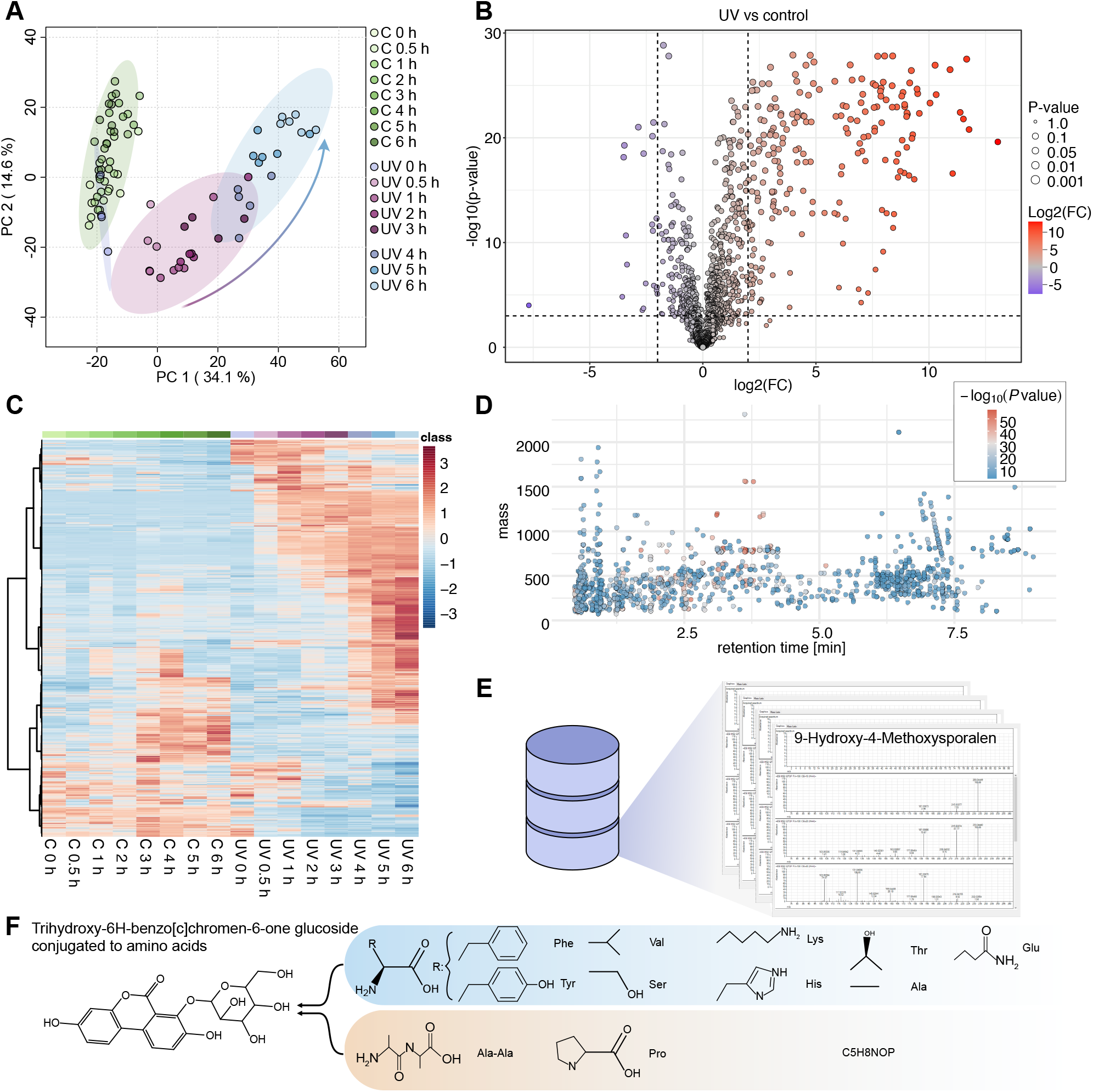
A chemodiverse time-course response to UV in the model zygnematophyte *Mesotaenium*. **(A)** Principal Component Analysis (PCA) of the features detected by untargeted metabolomic analysis by UHPLC QTOF MS. Principal Component (PC) 1 vs. PC 2, control samples (green) cluster together with UV 0 h samples (samples taken before UV-B irradiation was switched on). Samples during the first 3 h of UV-B exposure (pink) and samples after the exposure (blue) cluster together. Each timepoint has a specific shade of colour indicated on the right. The temporal dynamics of the progressive recovery are highlighy by an arrow. **(B)** Volcano plot of all UV-B treatment-derived data versus the control group of metabolomic samples x-axis: log2(FC) where Fold Change (FC) = |4.0|; y-axis: -log10(p) where the t-test p-value threshold is 0.001 false discovery rate (FDR). Grey: non-significant changes; red: enriched; blue: depleted. The volcano plot was generated by MetaboAnalyst ^111^, with equal group variance. **(C)** Metabolomic profiling. Each column is a timepoint, each row a feature. The data was auto-scaled, hierarchically clustered by ward and euclidian distance measure was applied. The colour gradient represents higher relative abundance in red and lower relative abundance in blue. The heatmap was generated by MetaboAnalyst^111^. **(D)** Scatter plot of features detected at a given mass over charge (m/z) and retention time (RT). **(E)** A metabolite database was created for *M. endlicherianum* SAG 12.97 with an example of MS/MS spectrum of a hydroxy methoxysporalen-like. **(F)** A *Mesotaenium*-specific compound, trihydroxy-benzo[c]chromen-one hexose was identified, which is conjugated to various amino acids.

236 features showed significant changes upon UV-B exposure; only 20 features showed depletion whereas 216 features showed an increase in abundance comparing all timepoints to control group (Figure 3B). Of these, 10 even increased by more than a thousand-fold (Figure 3B). As observed on differential gene expression changes, there is a time progression in the metabolite fingerprints. Features increased in abundance over time, and with time additional features contributed to the bouquet of compounds shaping the UV response (Figure 3C).

While research on streptophyte algae has immensely benefited from genomic and transcriptomic approaches^9^, with the first proteomic endeavors lined up^49^, our understanding of the diverse metabolites produced by zygnematophytes has largely focused on signaling molecules such as apocarotenoids^27,66^ and phytohormones^67^.

It is a topic of general interest in the field of early plant evolution to understand the evolutionary origin of phytohormone-based signaling cascades. Diverse phytohormones have been investigated, often leading to the realization that divergent signaling tracks or even ligands are likely involved in the molecular physiology of land plants’ closest algal relatives^44,67,68^. Ethylene stands in stark contrast to this. Endogenous ethylene appears to be produced in an aminocyclopropane-1-carboxylic acid (ACC)-dependent manner^69^ and exogenous application of ethylene triggers cell elongation as well as stress-associated gene expression patterns in Spirogyra^70^. Furthermore, Spirogyra genes homologous to proteins of the ethylene signaling cascade can fully or at least partially rescue the respective Arabidopsis knockout lines^69^. In our experiments, we found that a mass tentatively identified as the ethylene precursor ACC showed dynamic changes in abundance upon UV-B exposure: the signal intensity increased in a time-dependent manner by 1.5 times after 3 h and recovered swiftly after UV-B exposure ceased (Figure 4D). Indeed there are several homologs of the orthogroup^27^ that contains ETR1 that are upregulated (Figure 4D)—yet they do not contain the full ETR1 domain structure.

**Figure 4.**
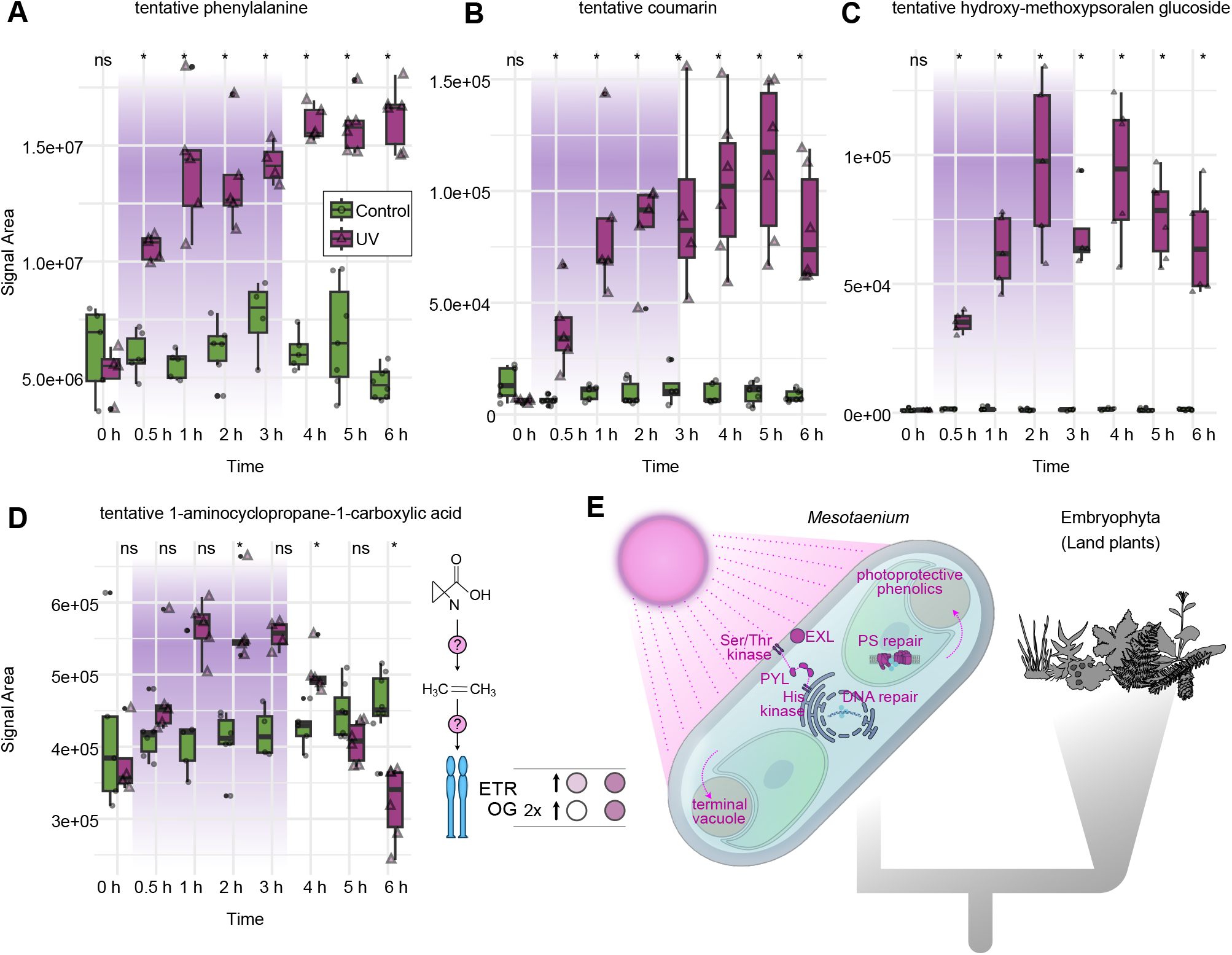
Temporally dynamic signature metabolites in the acclimation response to UV in *Mesotaenium*. A-D Box plots of four enriched metabolites upon UV-B. x-axis: time in hours; y-axis: signal area. Purple shading represents the time span during which UV-B irradiance occurred. Green: control, purple: UV-B samples. Significance levels shown on the top, *: p ≤ 0.05. (A) tentative phenylalanine; identified untargeted; (B) tentative coumarin; targeted; (C) tentative hydroxy methoxypsoralen glucoside; targeted; (D) tentative 1-aminocyclopropane-1-carboxylic acid; targeted. (E) Model of cell system response of *Mesotaenium* to UV-B stress. *Mesotaenium* as a representative of the closest algal relatives of land plants (Embryophytes) on a schematic phylogenetic tree. EXL: EXORDIUM-like, PYL: PYR1-like, PS: photosystem.

To move beyond analyses targeted at finding signaling compounds that are known from land plants, we here generated a comprehensive dataset of features with 1477 entities, at a high confidence level (ANOVA *P* value < 0.0001) for *Mesotaenium* as model system for the closest algal relatives of land plants (Figure 3D). An untargeted MS/MS–based profiling was performed to assign features of this *Mesotaenium* metabolite database, resulting in the annotation of 62 metabolites based in the manual interpretation of individual MS/MS spectra. They captured several compounds that can act as precursors for phenolics, phenolic compounds themselves, harboring derivatives which have yet not been described from other species (Figure 3E, suppl. Figure S4). *Mesotaenium* forms diverse combinations of conjugates derived from specialized metabolite core structures, resembling the added complexity also known from the phenylpropanoid pathway of different species of land plants^71-73^. A particularly noteworthy case is trihydroxy-benzo[c]chromen-6-one hexose, which we identified based on the well-described MS/MS spectra of the core structure^74^. While its abundance remains unchanged upon UV exposure, this compound represents a constitutive metabolite marker of *Mesotaenium*, detected in multiple conjugated forms with amino acids. (Figure 3F).

We thus next scrutinized specific compounds to understand the evolutionarily shared as well as unique biology that underpins the chemodiverse UV response in this model for the closest algal relatives of land plants.

### *Mesotaenium* produces specific phenolics under UV exposure

Most phenolics found in land plants derive from the shikimate pathway^71,75-80^. The aromatic amino acid phenylalanine is both the end product of the shikimate pathway and the entry point into phenylpropanoid biosynthesis. We found that within 30 minutes after UV-B exposure, phenylalanine levels nearly doubled (1.7-fold; P = 0.016) and steadily increased to a high of 3.4-fold as compared to control (*P* = 0.014; Figure 4A). This reflects not only that phenylalanine is an UV-B responsive amino acid, but it can be sourced into diverse phenol producing pathways downstream; this also opens up the possibility that its enrichment is owed to downstream steps of biosynthesis being blocked or hindered. Along these lines, we found diverse coumarin-like metabolites. Some coumarins have antioxidant properties and may act as reactive oxygen species (ROS) scavengers^81^. In embryophyte and algae, ROS triggered by UV-B exposure are a major culprit, damaging diverse biomolecules and (sub)cellular structures^82-84^. Some of these tentative coumarin derivatives increased by a factor of 6 already within 30 minutes, further increasing reaching a level of nearly 12-times the amount as compared to control levels within 5 h (Figure 4B).

A prominent UV-B-responsive metabolite was detected in *Mesotaenium* at RT 3.3 min in both ionization modes. In negative mode, the precursor ion at m/z 393.0826 (C_18_H_18_O_10_) showed a dominant neutral loss of 162.052 Da, consistent with cleavage of a hexose moiety, yielding an aglycone ion at m/z 231.0302. Subsequent fragment ions were consistent with sequential CO and CO_2_ losses from an aromatic lactone scaffold. In positive mode, the corresponding precursor at m/z 395.0979 displayed an analogous fragmentation series following hexose loss (m/z 233.0446 → 215.0340 → 187.0390 → 159.0443 → 131.0489), characteristic of conjugated benzopyranone/furanocoumarin-type chromophores^74^. The compound further exhibited UV absorbance maxima at 210, 310, and 340 nm, consistent with strong UV-absorbing aromatic systems.

To contextualize the aglycone structure, fragmentation patterns were compared with those of authentic furanocoumarin standards. An isopimpinellin (dimethoxypsoralen) standard showed the expected psoralen-type fragmentation series, and the *Mesotaenium* aglycone displayed closely related product ions, shifted by ∼14 Da relative to isopimpinellin, consistent with one fewer methyl equivalent. Additional qualitative similarity was observed when compared with published methoxsalen (8-methoxypsoralen) fragment ions. Although MS/MS does not allow assignment of substituent positions or sugar identity, the combined MS/MS behavior and UV–visible properties support the presence of a psoralen/furanocoumarin-like benzopyranone chromophore (Figure S4).

Among the compounds that showed the strongest UV-dependent kinetics was the furanocoumarin hydroxymethoxypsoralen-hexose, increasing upon UV-B exposure from a meagre baseline by nearly two orders of magnitude (95-fold) in abundance within 2 h after UV-B onset (Figure 4C). In land plants, furanocoumarins derive from the phenylpropanoid pathway-sourced coumarins^85-87^. The biosynthesis of furanocoumarins has - even in land plants - been subject to convergent evolution^88,89^. A key step towards furanocoumarins is the deoxygenation of *p*-coumaroyl-CoA, which has been found in diverse algae^90^, to the coumarin umbelliferone – the enzymes for this conversion have been recalcitrant to detection^89,91^. Thus, while the biosynthetic routes towards them remain obscure, tentatively assigned coumarins; it seems that coumarins and coumarin-derived compounds could be a feature of the UV response in *Mesotaenium*. This suggests shared conserved action of coumarins in land plants and their closest algal relatives; whether the coumarin-like compound detected here arises from an alternative biosynthetic pathway rather than the canonical land-plant phenylpropanoid pathway, such as a shikimate-derived or polyketide-like route, remains obscure.

Overall, our data suggest that *Mesotaenium* produces a wide range of different compounds. These include phenolics, several of which show a pronounced stress kinetic initiated by UV-B exposure. Importantly in all cases, the kinetics of compound accumulation progresses in the recovery phases, making them part of the acclimation response to UV in this zygnematophyte.

### On the diverse routes of evolution of integrated stress responses

Land plants are characterized by their chemodiverse specialized metabolism. Here, phenolics are of key importance. Embryophytic phenolics sourced from the phenylpropanoid pathway are both key specialized metabolites in response to terrestrial stressors but also their structure, growth, and development^79,92^. Our data point to coumarin-like derivatives differentially accumulating upon UV exposure. Likely, enzymes homologous to those integrated in the embryophytic phenylproanoid pathway are acting in zygnematophytes—even though they lack homologs for parts of the land plant canonical pathway enzymes. This makes it doubtful that the phenylpronpaid pathway acts in zygnematophytes. Instead, the homologous enzymes have likely been rewired, assembling a different pathways^13,93^. Indeed, the enzyme families involved in the production of phenolics radiated and diverged during repeatedly evolution^58^; high evolutionary versatility indicated by these radiations suggests functional divergence of the pivotal enzymes involved in this pathway^58^. It thus is one of the readouts in the layered response system to stress acting in embryophtes^2^, of which homologous sub-networks occur in their closest algal relatives^18,26,27,31,93-95^. From a genetic point of view, the function of many of these genes remains to be elucidated, their presence depicts the ancestral gene pool from which the embryophytic stress response genes emerged^1,96^. And genetic approaches often move from perception, to transduction, to output. If we take a physicochemical standpoint, we can flip this rationale: upon application of the stress, we observe the output (for whose detection universally applicable omic techniques are particularly apt) and work our way to the regulatory level.

### A model for systems acclimation to UV in zygnematophyte cells

UV radiation (UVR) is an abiotic stressor especially important in terrestrial habitats. In aquatic ecosystems, the overlying water column protects the sublittoral algae against most of the radiation. Even though UVR penetrates pure water deeper compared to other radiation, freshwater usually teems with other organisms and dissolved compounds, such as phenolics, which filter and absorb the UVR efficiently^97^. UV-B radiation can directly induce alterations of biomolecules like DNA, proteins and chloroplasts or indirectly via metabolically toxic ROS^82,98^. There are different mechanisms to remedy or evade such damage. These mechanisms include DNA repair processes, UVR avoidance, the biosynthesis of antioxidants and UV-screening compounds^98^. UV-screening compounds may absorb the energy of UVR, acting as mere shields, disposing of the previously harmful energy in form of heat, thereby preventing damage to essential biomolecules^82^. The absence of such mechanisms would result in harmful effects, such as reduced or inhibited growth^99,100^, reproduction, survival^101^, protein stability in photosystem II (PSII), pigmentation^102^, and activity of the key enzyme for CO_2_ fixation, Ribulose-1,5-bisphosphate carboxylase-oxygenase (RuBisCO)^103^.

Terminal vacuoles in zygnematophytes, such as those characterised for the unicell *Closterium moniliferum* are part of the cell biological acclimation responses to adverse environmental cues^104,105^. In these terminal, or polar vacuoles, precipitates of barite (BaSO_4_), celestite (SrSO_4_), strontium citrate, and other salts in form of crystals are commonly found^106,107^; further, the filamentous zygnematophyte *Zygnema* is known to store phenolic compounds in its vacuoles^107^. These occurred most notably during abiotic stress such as UVR^6^. The observed compounds encompassed glycosylated, highly branched phenolics in complex with iron Fe^3+^, resulting in a purple colour. These pigments serve as a sunscreen as well as reduce the risk of excessive iron inducing oxidative stress due to the Fenton reaction^108^. In another unicellular zygnematophyte, *Ancylonema alaskanum*, vacuoles with brownish pigments, putatively polymerised Fe-complexes of purpurogallin, were observed upon UV and VIS irradiation^109^. It is conceivable that the here discussed glycosylation of coumarin-like metabolites may enhance their solubility and transport to the vacuole, which could serve as a disposal site after ROS salvaging. Interestingly, flavone glycosides accumulation is known in *Marchantia polymorpha*^19^, which may absorb UV-B as well as scavenge ROS.

Altogether, we here identified a systems response that likely includes the action of conserved kinase-based stress nexuses, photoreceptors, plastid biology homeostats, specialized metabolites, and cellular alterations in cell wall and vacuoles (Figure 4E). Homologs of ETR1, which were found to be a central node in stress signalling, likely predates the divergence of Embryophytes and their closest algal relatives^27^. Such conserved histidine kinases likely mediate the stress response, connecting to Ser/Thr hubs, further modifying the downstream growth and stress effectors like the previously mentioned EXL, forming a kinase backbone coupling perception to acclimation and adaptation. Photoreceptors like UVR8 detect incident UV-B radiation that, in land plants, activate the downstream signalling cascade to induce acclimation. Here we observed many such downstream acclimation responses, including the regulation of genes involved in DNA and protein repair mechanisms, photosynthesis and cell shape adaptation via e.g. *EXL* and changes in the metabolite fingerprints that speak of phenolic production.

Concertedly, our observations describe a model for UV-B acclimation responses in zygnematophytes from cue perception, signaling, physiological and cell biological alterations, up to the accumulation of compounds in UV warding (Figure 4E). Some of these were likely part of the plant terrestrialization toolkit; that said, many responses are likely specific to the closest algal relatives of land plants—with the bouquet of phenolics being a point in case. Only a phylodiverse approach that synthesises all data will allow to extract the shared core repertoire of the earliest land plants.

## Conclusion

The origin of land plants is a complex question that requires the synthesis of diverse avenues^110^. One avenue is an experimental approach in the closest algal relatives of land plants: the zygnematophytes. Often unicellular, zygnematophyte cells contextualized with land plants cell biology allow for a synthesis of the logic of stress responses^49,93^. Here, we synthesized a model for UV response in zygnematophyte cells (Figure 4E), shedding light on the integrated molecular physiological toolkit to a prime stressor on land: UV irradiance.

## Materials availability

This study did not generate new, unique reagents.

### Data and code availability

- All RNA sequencing (RNA-seq) data generated in this study have been deposited in the NCBI under BioProject accession number BioProject: PRJNA1266594.
- All metabolomic files generated in this study are available at Metabolights public repository under the identifier MTBLS8088.

## Acknowledgement

We thank Prof. T. Friedl from the Department of Experimental Phycology and Culture Collection of Algae (SAG), Prof. S. Neugart and René Heise at the University of Göttingen for their support. J.d.V. is grateful for funding by the German Research Foundation (DFG) grants 509535047 (VR 132/10-1) and 569599232 (VR 132/18-1) and the grants 440231723 (VR 132/4-1), 528076711 (VR 132/13-1) within the framework of the Priority Programme “MAdLand – Molecular Adaptation to Land: Plant Evolution to Change” (SPP 2237). J.d.V. further thanks the European Research Council for funding under the European Union’s Horizon 2020 research and innovation programme (Grant Agreement No. 852725; ERC-StG “TerreStriAL”) and the Horizon Europe programme (Grant Agreement No. 101230161; ERC-CoG “StreptoProgram”). I.F. acknowledges funding by the DFG grants 495720893 (INST 186/1434-1) and 440232164 (FE 446/14-1) within the framework of the Priority Programme “MAdLand – Molecular Adaptation to Land: Plant Evolution to Change” (SPP 2237). C.F.K is grateful for being supported through the International Max Planck Research School (IMPRS) for Genome Science within the framework of the ‘Göttingen Graduate Center for Neurosciences, Biophysics, and Molecular Biosciences’ (GGNB) at the University of Goettingen.

## AUTHOR CONTRIBUTIONS

Conceptualization, C.F.K., J.d.V.; data curation, C.F.K., I.N.A., and K.F.; investigation, C.F.K., and J.d.V.; experimental design, C.F.K., I.N.A., J.F.J. and J.d.V; experimental work, C.F.K., I.N.A., T.D., M.L, and J. d. V.; resources, I.F. and J.d.V.; writing – original draft, C.F.K., I.N.A. and J.d.V.; writing – review and editing, all authors; visualization, C.F.K. and J. d.V.; funding acquisition, J.d.V.

## METHODS

### RESOURCE TABLE

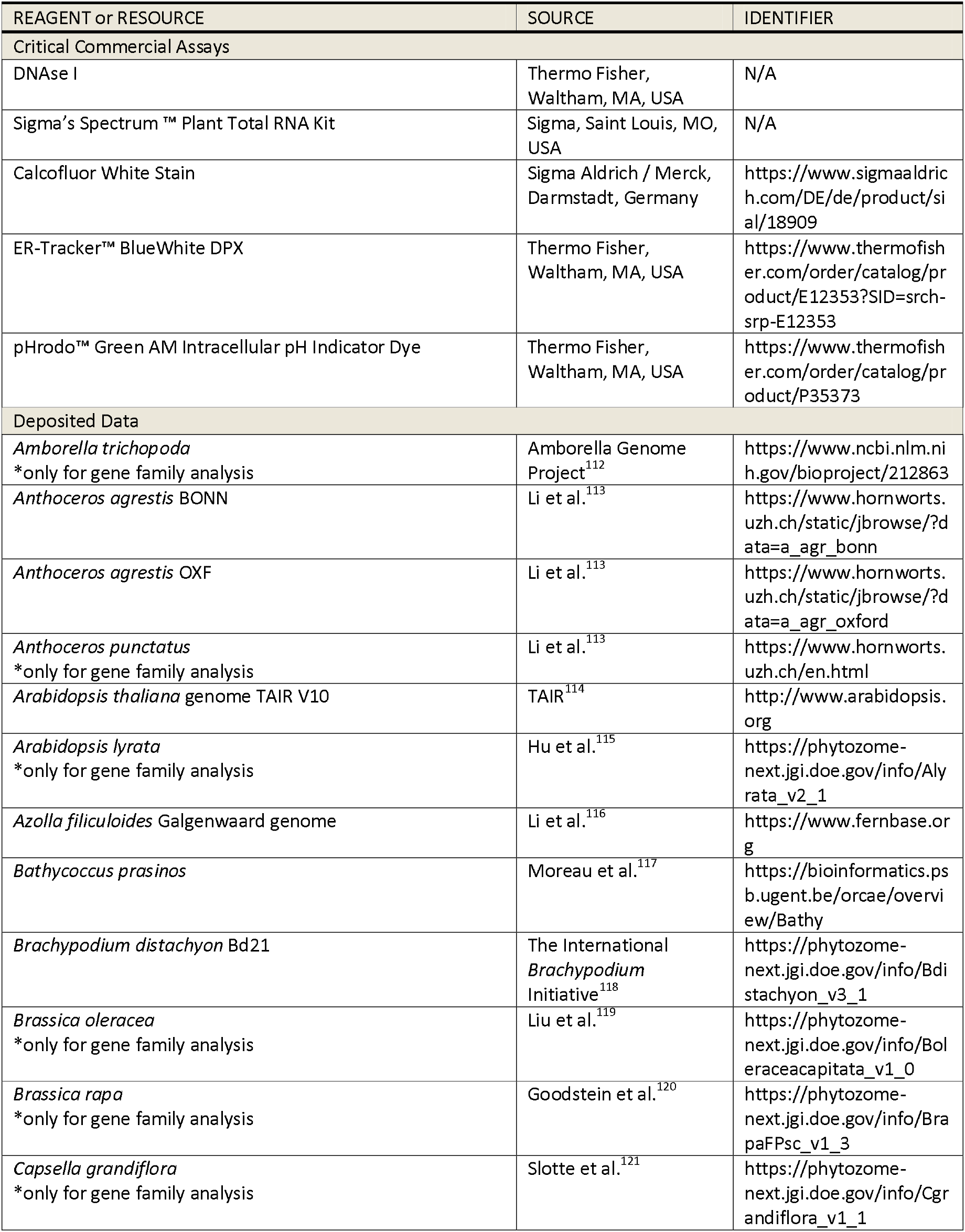
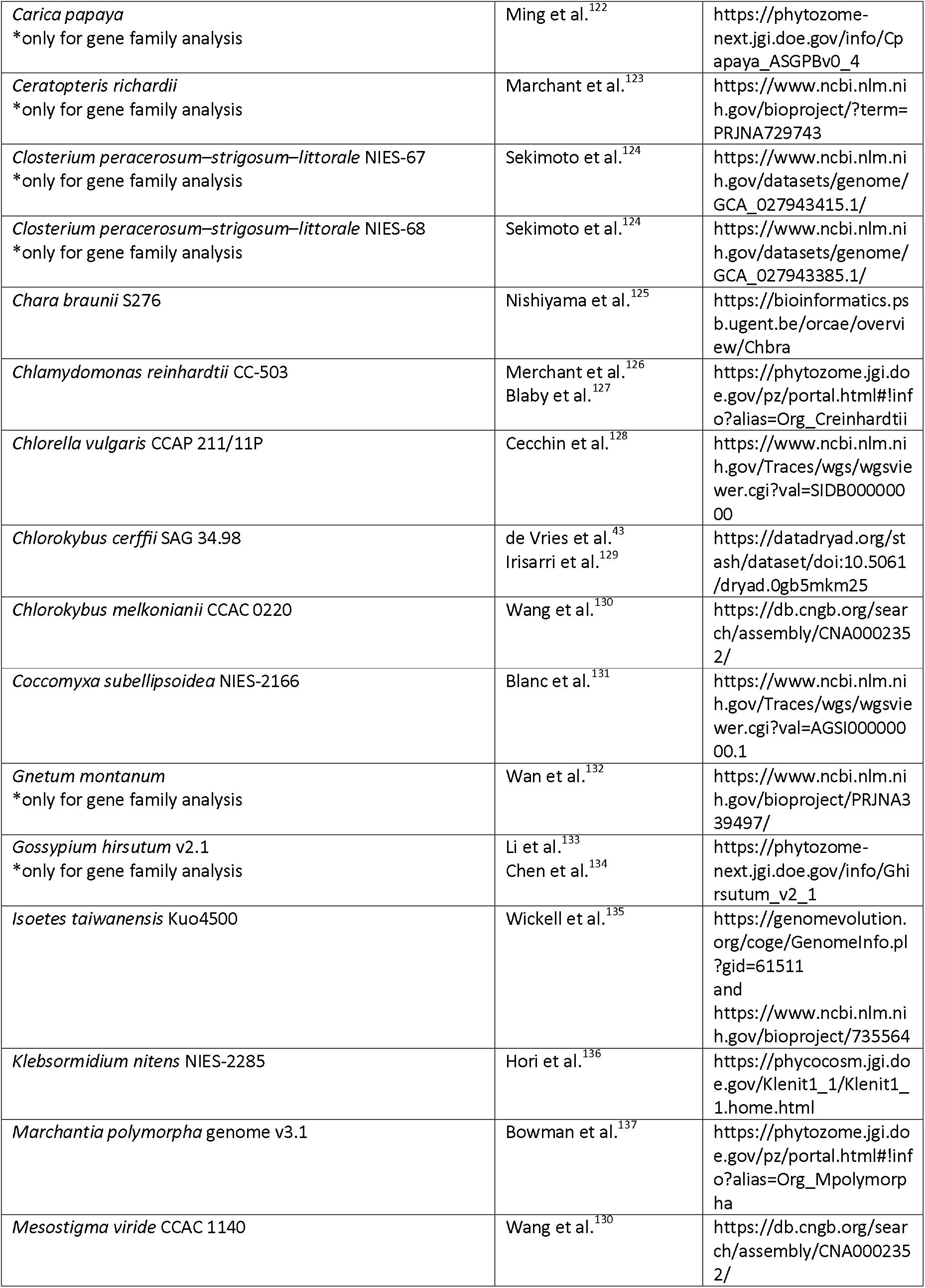
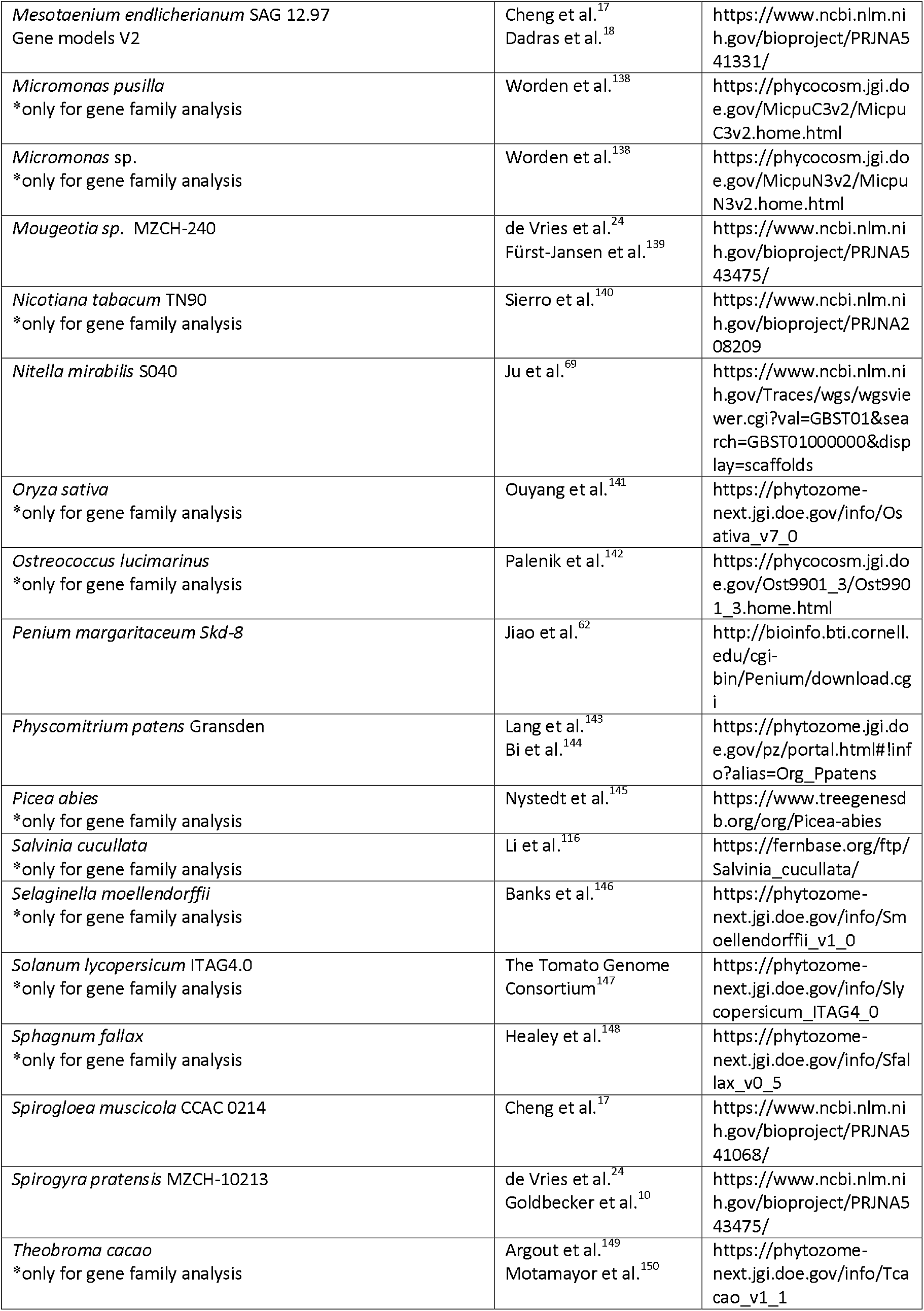
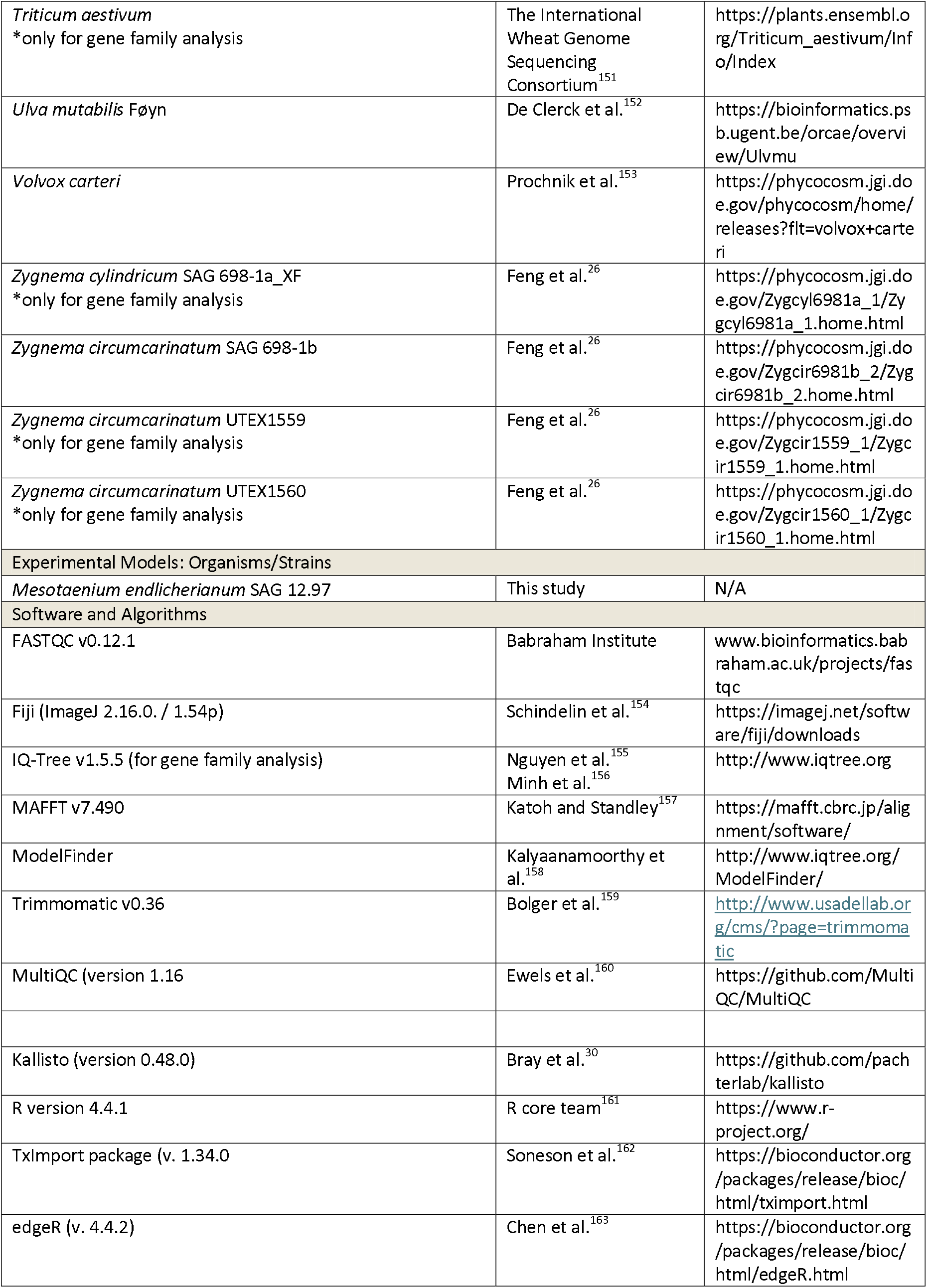
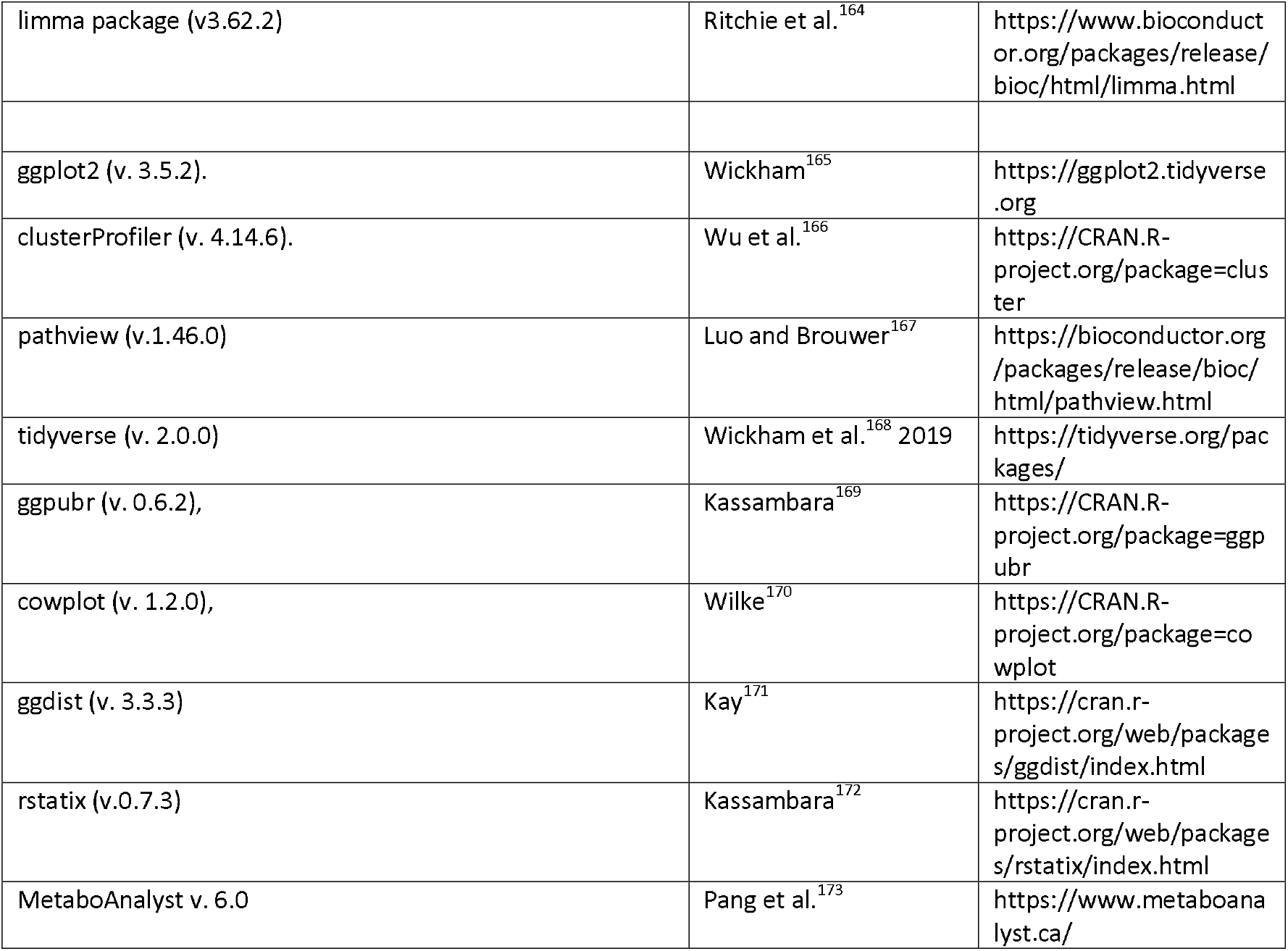

### EXPERIMENTAL MODEL AND SUBJECT DETAILS

#### Algal strains

*Mesotaenium endlicherianum* SAG 12.97 was obtained from the Department Experimental Phycology and Culture Collection of Algae at Göttingen University (EPSAG)^174^.

## METHOD DETAILS

### Cultivation of Algae on Solid Medium

For stock culturing, *M. endlicherianum* was stored on solid Bold’s Basal Medium (BBM) with vitamins and triple nitrate (3NBBM+ V) according to the recipe of Schlösser et al.^175^ (see also Bischoff and Bold ^176^ and Starr and Zeikus ^177^) at 15 ºC under full-spectrum fluorescent lamps (5-10 μmol photons m^-2^ s^-1^; constant light). Stock cultures between 21 and 28 days were used for the inoculation of experimental plates.

For easier sampling and reduction of agar and media contamination in metabolite samples, the agar was covered with cellophane (folia®, Max Bringmann KG). The cellophane was cut to petri dish size, autoclaved and applied using a heated Drigalsky spatula and five drops of sterile ddH_2_O. The biomass of one stock plate was diluted in 10 mL liquid medium at room temperature and 300 μL of the solution was spread on each plate using a sterile cooled Drigalsky spatula. Plating occurred under the sterile bench with sterile technique. The plates were closed using 3M Medical Micropore Tape, as it was shown to ensure better CO_2_ exchange than parafilm ^178^.

For the UV-B exposure experiments, the plates were grown at 80 - 90 μmol photons m^-2^ s^-1^ (Niello® LED 300W, 380–740 nm spectrum; Supplementary Fig. S1A) for seven days with a 16/8 h light/dark cycle, before being transferred to the UV setup location for 24 h acclimation before the experiments.

### Experimental Set-Up

The UV setup used the same grow lights as previously described in addition to two UV-B lamps used in conjunction, the UV-B Broadband TL fluorescent tube lamp with a UV-B spectrum of 290 to 315 nm, and the UV-B Narrowband TL fluorescent tube lamp with a UV-B spectrum between 305 and 315 nm, which peaks at 311 nm (lamp wattage: 40 W and 20 W; length: 1213.6 mm and 604 mm respectively, Philips, the Netherlands; spectrum Supplementary Fig. S1A). They produced a UV-B intensity ranging from 1.7 - 2.9 W m^−2^, depending on location of the individual petri dish. The photosynthetically active radiation (PAR) values were 40 - 50 μmol photons m^-2^ s^-1^and 50 - 60 μmol photons m^-2^ s^-1^ for UV-B and control, respectively. While PAR was applied with the previous 16/8 h light/dark cycle, for UV treated samples UVR was applied for 3 h (Fig. X) To reduce condensation on the lids of the petri dishes, fans were installed to create air circulation. Temperature and humidity were monitored using a Tempo Disc™ (BlueMaestro). The temperature and humidity ranged from 17.2 ºC to 20.6 °C with an average of 19.2 ºC and 29.7 % to 51.3 % with an average of 40.4 %, respectively.

10 min before the experiment started, the micropore tape around all the plates and the lid were removed, to avoid UV-B irradiation being filtered out by the plastic lid, as contamination within the following max. 6 h can be neglected. For the multiple day experiment, the lids were left on, which resulted in a reduced UV-B intensity of 0.6 – 1.2 W m^-2^. The measurements of photosystem II (PSII) maximum efficiency (Fv’/Fm’) were taken at three independent locations on two plates. Morphology assessment through microscopy was performed at 0 h, 3 h and 6 h using the Olympus BX-60 microscope (Olympus, Japan) with DIC equipped with JENOPTIK GRYPHAX® PROKYON camera and JENOPTIK GRYPHAX® microscope camera software (version V2.2.0) (JENOPTIK AG, Jena, Germany). The imaging occurred immediately after measurement. Additional confocal images were taken after 6 h of constant UV-light as well as after 6 h of 3 h UV and 3 h recovery. The cells were stained using 1% Calcofluor White Stain (Sigma-Aldrich, Germany), Invitrogen™ pHrodo™ Green AM Intracellular pH Indicator Dye and Invitrogen™ ER-Tracker™ Blue-White DPX. The images were taken using the Zeiss LSM 980 Confocal Laser Scanning Microscope equipped with diode lasers 405nm, 445, 488, 514nm; diode-pumped solid state lasers 543, 561, 594, 639nm all with the objectives 40x/1,4Plan-Apochromat and an Axioxam 512 color camera using Zeiss Zen blue software V3.2 (Zeiss, Jena, Germany). Sampling was performed as follows: two plates at each time point of the experiment were taken out of both settings, *Mesotaenium* was removed from the cellophane and 1/4 of each plate pooled (depending on setting) for transcriptomics and 3/4 of each plate pooled for metabolite analysis. Pooling was done to balance the varying irradiation intensity. Samples were frozen immediately using liquid N_2_ and stored at -70 °C, to stop any (non-) enzymatic reaction which would distort the results. The experiments were repeated three times to obtain biological triplicates.

### RNA isolation and sequencing

The Sigma Aldrich Spectrum™ Plant Total RNA-Kit was used for RNA extraction, following the “Spectrum Plant total RNA Isolation protocol A” protocol with the following modifications. 500 µL of the prepared lysis solution/2-mercaptoethanol mixture was added to the sample along with 1/3 beads from MPI Lysing Matrix E. This was immediately vortexed and the biomass homogenized using a Biospec Mini-BeadBeater for 30 s at 5000 rpm. Afterwards, 200 µL lysis solution/2-mercaptoethanol mixture was added, samples placed in an ultrasonic bath (DK Sonic DK-300H, 40 KHz, 120 W) for 2 min with constant ultrasonication. After incubation at 56 ºC for 5 min, the solution was transferred to a new Eppendorf tube, avoiding the BeadBeater beads and centrifuged for 1 min at 16 000 x g to remove cell debris and remaining beads. The supernatant was loaded onto the filtration columns. To bind RNA, protocol A with increased amount of Binding Solution of 750 µL was used and subsequently, the manufacturers protocol was followed. RNA concentration and quality was assessed using a NanoDrop Spectrophotometer ND-1000 and 1% agarose gel. 1 µL 6 x loading dye was added to 4 µL sample and incubated at 60 ºC and on ice for 5 min each before loading onto the gel. The remaining sample was treated with DNAse I according to manufacturer’s protocol (Thermo Fisher). The sequencing was performed at Novogene (Cambridge, UK) on an Illumina NovaSeq 6000 platform, generating 150 bp long paired reads.

The following 50 and 30 sequence adaptors were used: 50-AGATCGGAAGAGCGTCGTGTAGGGAAAGAGTGTAGATCTCGGTGGTCGCCGTATCATT-30 (50 adaptor) and 50-GATCGGAAGAGCACACGTCTGAACTCCAGTCACGGATGACTATCTCGTATGCCGTCTTCTGCTTG-30 (30 adaptor)

### Transcriptome analysis

The quality of the raw and processed reads was checked using FastQC (version 0.12.1) ^179^ and MultiQC (version 1.16) ^160^. Trimmomatic (version 0.39) ^159^ was used to process raw reads with the setting “ILLUMINACLIP:adapters/adapter.fasta:2:30:10:2:True LEADING:26 TRAILING:26 SLIDINGWINDOW:4:20 MINLEN:36”. The processed reads were pseudoaligned to cDNA of *M. endlicherianum* SAG 12.97 gene model V2 ^18^ with Kallisto (version 0.48.0) ^30^ with 100 bootstraps. The samples were sequenced with the rf-stranded technique. Subsequent analysis was performed using R version 4.4.1. The transcript-level estimates were imported via the TxImport package (v. 1.34.0) ^162^ and estimated counts calculated based on abundance with “lengthScaledTPM”. edgeR (v. 4.4.2) ^180^ was used to filter (omitted genes with less than 1 count per million in at least 3 samples) and normalise (upperquartile method ^181^) the counts. The data was subsequently transformed to log2-count per million via the voom function in the limma package (v3.62.2) ^182,183^. The principal component analysis was performed with stats (v. 4.4.1) and visualised with ggplot2 (v. 3.5.2)^165^. Functional annotation of genes via eggnog-mapper retrieved from Dadras et al. 2023 a. Gene ontology (GO) over-representation analysis (ORA) was performed using the clusterProfiler enricher function (v. 4.14.6)^166^ using a q-value and adjusted p-value cutoff of 0.05 each. We used pathview^167^ (v.1.46.0) and KAAS^184^ (KEGG Automatic Annotation Server) for additional information on the transcripts.

The p-values were calculated using the Student test (Student^185^) and adjusted with the Benjamini and Hochberg (BH; Benjamini & Hochberg^186^) method.

### Sample Preparation and Metabolite Extraction

Cultures were transferred into Eppendorf tubes and lyophilized overnight. Two tungsten beads were added to each tube, and the material was pulverized using a Mixer Ball Mill MM400 (Retsch). Approximately 5 mg of dried and homogenized algal material was extracted with 500 μL of 70% (v/v) methanol, following the protocol described by Kelly et al.^187^. After centrifugation, 200 μL of the supernatant was transferred to an LC vial and dried using a SpeedVac. Prior to analysis, samples were reconstituted in 20 μL methanol and subsequently diluted with 60 μL Milli-Q H_2_O.

### Untargeted Metabolomics Analysis of *Mesotaenium* Cells

Samples were analyzed using an ultra-high-performance liquid chromatography (UHPLC) system (1290 Infinity, Agilent Technologies, Santa Clara, CA, USA) coupled to a quadrupole time-of-flight mass spectrometer (QTOF-MS; 6546 UHD Accurate-Mass QTOF, Agilent Technologies). A volume of 2 μL was injected onto an Acquity HSS T3 column (1.8 μm, 2.1 × 100 mm; Waters Corporation, Milford, USA) maintained at 40 °C. Metabolites were eluted at a flow rate of 0.5 mL min^-1^ using a binary gradient of water (0.1% formic acid) and acetonitrile (0.1% formic acid), as described by Feussner et al. ^188^. High-resolution accurate mass mass spectra were acquired over a mass range of m/z 80– 1700 using an electrospray ionization source operated in both positive and negative ionization modes. Raw MS data were processed using MassHunter Profinder B.08.00 (Agilent Technologies) employing recursive feature extraction, as described by ^189^. The resulting dataset was normalized to sample dry weight and subjected to statistical analysis using MetaboAnalyst 6.0 ^111^.

All MS files generated in this study are available at Metabolights public repository ^190^ under the identifier MTBLS8088.

### MS/MS-Based Metabolite Annotation

Additional Auto-MS/MS and All-Ions MS analyses were performed using collision energies ranging from 10 to 40 V to support systematic annotation of phenolic metabolites in *Mesotaenium* extracts. Metabolite annotation was conducted as follows: molecular formulas of precursor and fragment ions were calculated based on monoisotopic mass; diagnostic fragments were identified through searches in METLIN, KEGG, and PubChem databases, as well as available literature; conjugated metabolite forms were manually predicted based on MS/MS spectral interpretation, considering neutral losses, retention times, and ionization modes and All MS/MS spectra were imported into an in-house compound database and library (PCDL; Meso_database).

### Controls

Parallel extraction and LC–MS analysis of uninoculated agar powder were performed as a negative control to account for background metabolites originating from the growth medium. A dedicated LC column was used to minimize carryover and cross-contamination.

### QUANTIFICATION AND STATISTICAL ANALYSIS

For all statistical analysis R (v 4.4.1)^161^ was used. All p-values were adjusted for multiple testing via Benjamini–Hochberg procedure^186^.

For the analysis of Fv’/Fm’ (Fig. 1c), the packages tidyverse (v. 2.0.0), ggpubr (v. 0.6.2), cowplot (v. 1.2.0), ggdist (v. 3.3.3), and rstatix (v.0.7.3) were used (see Resource table). Normality was assed via the Shapiro-Wilk test^191^. As not all values meet the assumption of normality, non-parametric Wilcoxon rank-sum^192^ test (Mann-Whitney U test^193^) was employed. The data was visualized and quadratic polynomial trend lines were fitted to each treatment phase (control, UV stress, UV recovery) using least-squares regression.

All micrographs show representative images.

For the analysis of transcriptomics data (Fig. 2a-d), the p-values were calculated using the Student test^185^. The change of transcription was assessed via a regression (suppl. Fig.) where the statistical significance of the slope differing from 1 (perfect persistence) was evaluated using a t-test and a two-sided p-value was computed to test whether early changes significantly deviated from perfect persistence. Pearson’s correlation coefficient was calculated to quantify the overall association between log2FC at 0.5 h and 2 h, MetaboAnalyst v. 6.0 was used for statistical analysis and visualisation of metabolomic response (Fig. 3a-c). Statistical Analysis [one factor] was selected, a plain text file with samples in columns (unpaired) and peak intensities submitted. Data was auto scaled. False discovery rate (FDR) was always set to 0.001, fold change to 4.0.

The trend of specific feature abundance development over the time course of the experiment (Fig. 4A-D) was analysed and visualised using the rstatix package (v.0.7.3). Normality was assed via the Shapiro-Wilk test^191^. As not all values meet the assumption of normality, non-parametric Wilcoxon rank-sum test^192^ (Mann-Whitney U test^193^) was employed.

