## Supplementary Fig. for "Chemodiverse cell systems responses to UV in an algal sister of land plants"

### Supplement to “Chemodiverse cell systems responses to UV in a closest algal relative of land plants”

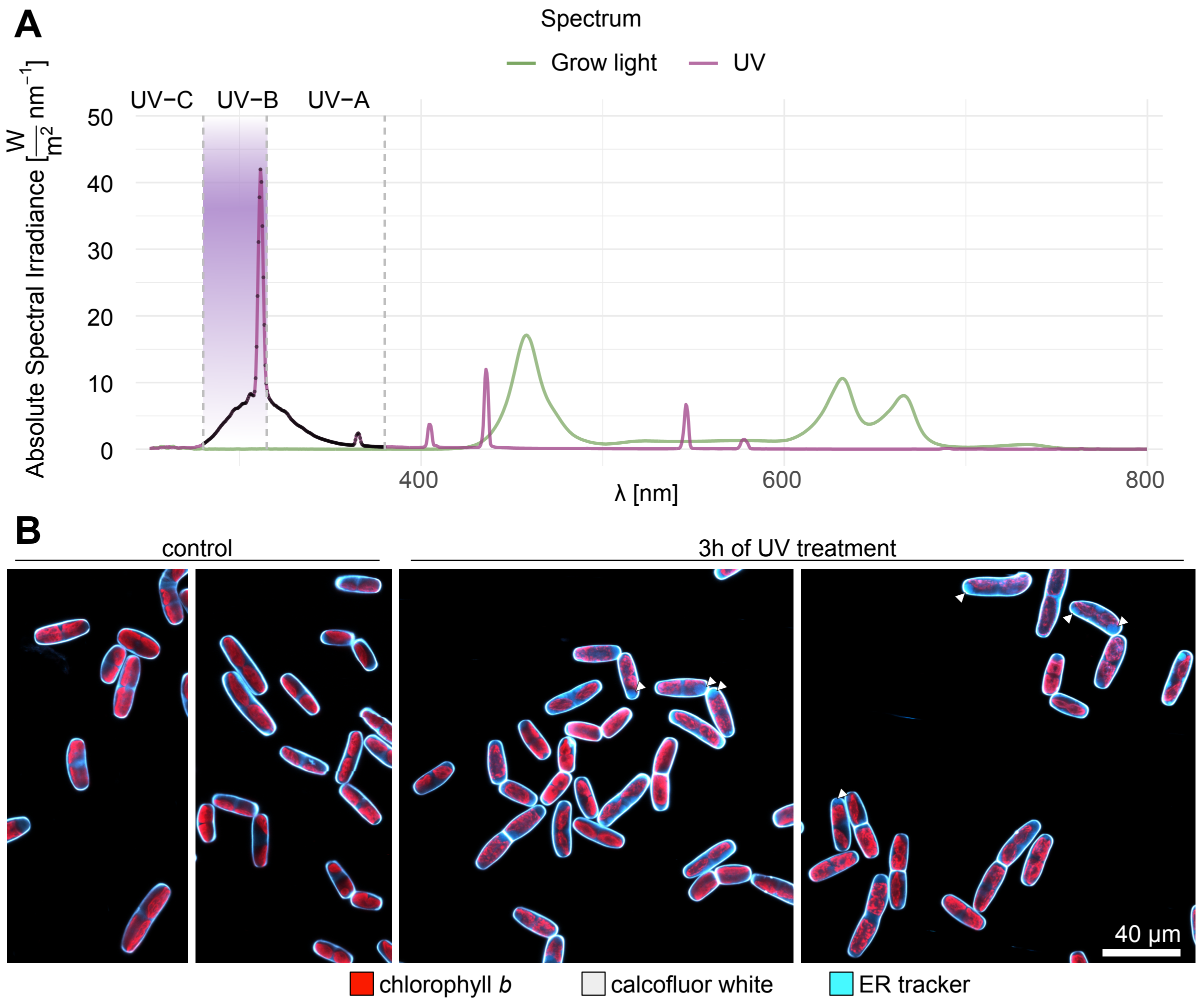


**Figure S1, related to Figure 1. (A)** Light Spectrum. Absolute spectral irradiance [W/m$^2$/nm] for each wavelength (λ [nm]). Yellow depicts the spectrum of the grow lights used, purple the spectrum of the UV-B lamps used. Grey vertical dashed lines along with denotation of UV-C, UV-B, UV-A on the top depict the according wavelength boundaries. Purple background indicates the wavelengths classified as UV-B. **(B)** Confocal microscope panel of control and 3 h continuous UV-B exposure. Chlorophyll autofluorescence (red) was used as well as stains for cell wall (calcofluor white, grey), and ER tracker (blue). Scale indicated in bottom right corner of each image.

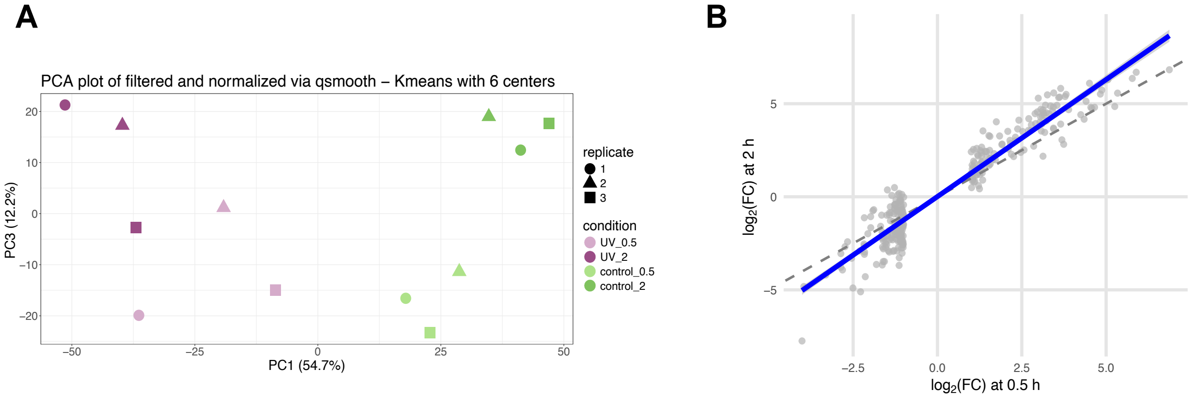


**Figure S2, related to Figure 2: (A)** PCA plot of filtered and normalized via qsmooth – Kmeans with 6 centers; PC1 versus PC3, shape indicating replicate, color condition. **(B)** Regression. $\log_2$(FC) at 0.5 h vs $\log_2$(FC) at 2 h of differentially expressed genes. Grey dashed line represents perfect persistence, blue line represents the observed trend.

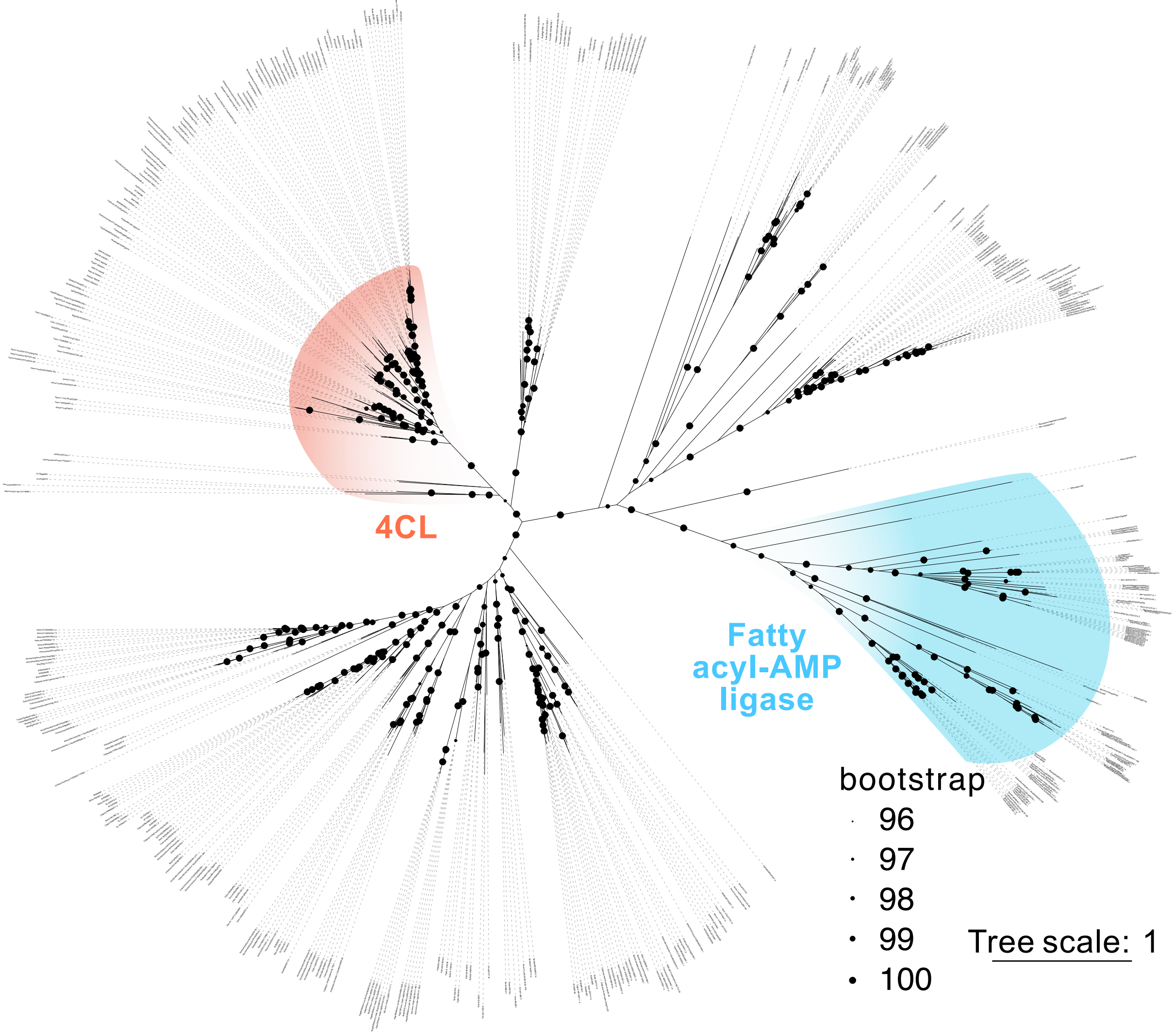


**Figure S3, related to Figure 2:** Full phylogeny shown in Figure 2f, including all genome names.

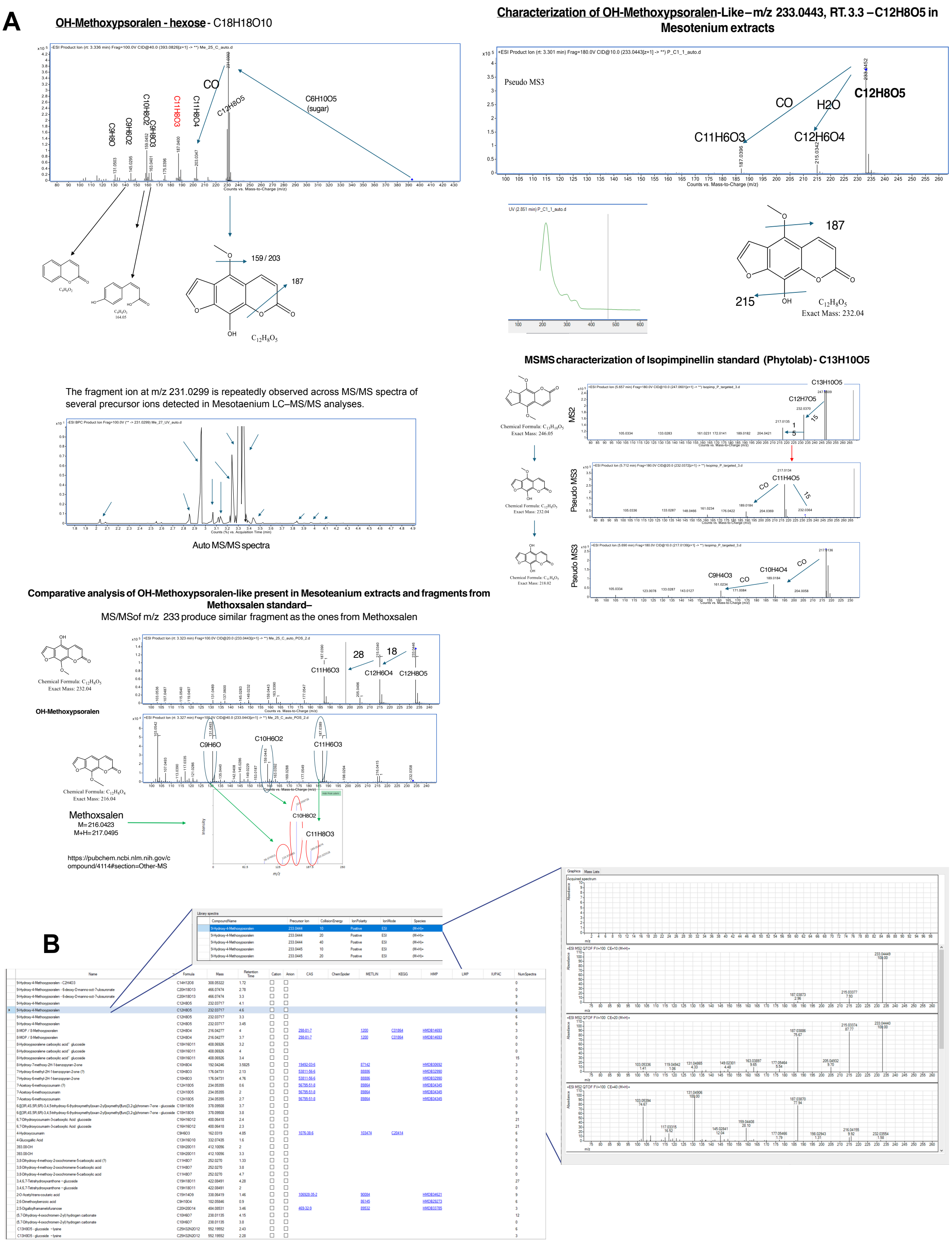


**Figure S4, related to Figures 3 and 4. (A)** Metabolite identification workflow: i) Selection of abundant ion (m/z 393.0823) and fragmentation by MSMS; ii) Manual inspection of MSMS spectra, where the ion formula ($C_{18}H_{18}O_{10}$) and its neutral loss with the fragment $C_{12}H_8O_5$ shows a loss of sugar ($C_6H_{10}O_5$); iii) the absorption at 280 and 310 nm are compatible with coumarins; iv) annotation of 9-hydroxy-4-methoxysporalen 9 glucoside. **(B)** Screenshots of MesotaeniumDB database, showing i) a list of compounds with their formula, mass, retention times, and, where present, CAS, ChemSpider, METLIN, KEGG, HMP, LMP and IUPAC identifiers; ii) Library of spectra for selected compound, with precursor ion, collision energy, ion polarity, ion mode and species; iii) m/z spectra.
